# Delayed Extracellular Matrix Negative Feedback Contributes to Nascent Adhesion Dynamics: Mathematical Modeling and Analysis

**DOI:** 10.1101/2025.05.06.652413

**Authors:** Vivi Andasari, S. Syafiie

## Abstract

Cell migration, both *in vivo* and *in vitro*, is a complex process governed by mechano-chemical interactions between cells and the extracellular matrix (ECM). These interactions, mediated by cell membrane receptors called integrins, involve bidirectional signaling between the intracellular actin cytoskeleton and the ECM. Integrins bind with cytoplasmic proteins to form adhesion complexes of varying sizes and maturity, which play crucial roles in cellular processes such as cell migration. Among these complexes are nascent adhesions–the smallest and earliest observable structures that emerge within the lamellipodium and are associated with rapid cell motility. While other adhesion types have been extensively studied, the mechanisms regulating nascent adhesions remain poorly understood.

Here, we develop a mathematical model describing the bidirectional signaling between the actin cytoskeleton and ECM that controls nascent adhesion dynamics. Our framework employs a system of delay differential equations to capture the temporal coupling between actin polymerization-driven adhesion formation and force-dependent ECM displacement. The model demonstrates that nascent adhesions, initiated by actin polymerization, exert forces on the ECM, whose delayed displacement provides negative feedback that limits adhesion growth. Numerical simulations reveal that this delayed ECM feedback mechanism reproduces the characteristic lifetime and dynamics of nascent adhesions, with quantitative agreement with experimental observations. Our results suggest that delayed ECM negative feedback is a key regulator of nascent adhesion turnover, providing new insights into the spatiotemporal control of cell migration.

## 1 Introduction

Cell migration, whether on 2D substrates or within 3D interstitial extracellular matrices (ECMs), relies on adhesive migration mechanisms involving a combination of adhesion and de-adhesion events orchestrated by various cellular strategies Friedl et al. [1998]. Cell adhesions to the ECM are complex interactions between the intracellular actin cytoskeleton (inside of the cell) and ECM components (outside of the cell) that are mediated by cell surface receptors called integrins, which are connected to associated proteins in the cytoplasm Hotchin and Hall [1995], Zamir and Geiger [2001], as schematically illustrated in Fig. 1. These interactions involve bi-directional signaling, from the intracellular cytoplasm to the extracellular environments (inside-out) and vice versa (outside-in) Berrier and Yamada [2007]. The molecular interactions between the intracellular and extracellular environments via cell-matrix adhesions are intricate and highly regulated both spatially and temporally Wiseman et al. [2004].

**Figure 1:**
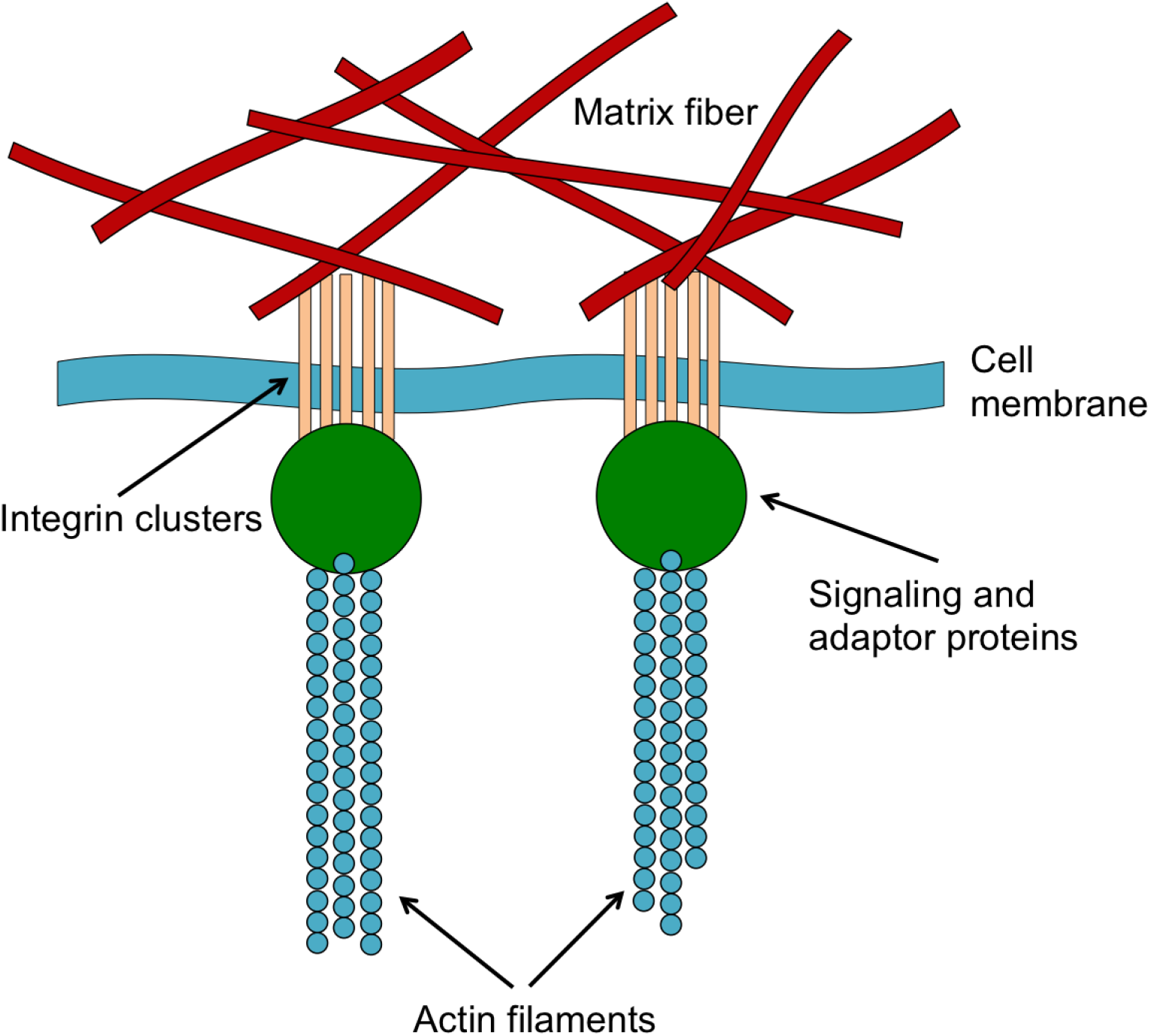
Generalized schematic diagram of nascent adhesion.

Integrin-based adhesions exhibit a variety of sizes, morphologies, and locations, depending on the cell type and its surrounding environment. In cells migrating on a two-dimensional (2D) substrate, these adhesions have been categorized into nascent adhesions, focal complexes, focal adhesions, and fibrillar adhesions Lock et al. [2008], Huttenlocher and Horwitz [2011], all representing distinct stages of adhesion maturation. Nascent adhesions are newly formed, small (*<* 0.25 *µ*m in diameter), dynamically transient, and dot-like structures. They contain a smaller repertoire of proteins compared to larger adhesions. Typically, integrin *β*_1_, kindlin, talin, vinculin, *α*-actinin, and paxillin are found within nascent adhesion structures Gardel et al. [2010], Bachir et al. [2014]. Focal complexes are approximately *∼*0.5 *µ*m in size and contain integrins, phospho-paxillin, FAK, *α*-actinin, talin, and myosin II, whereas focal adhesions are considerably larger, with an elongated size of 1 − 5 *µ*m, and comprise integrins, paxillin, FAK, talin, zyxin, vinculin, VASP, and myosin II. Fibrillar adhesions, the largest adhesion structures (*>* 5 *µ*m) are enriched with integrins, dephospho-paxillin, FAK, talin, vinculin, VASP, *α*-actinin, and tensin Choi et al. [2008], Gardel et al. [2010]. Nascent adhesions undergo rapid assembly and disassembly (turnover) within 60 seconds Choi et al. [2008]. They continuously appear at the leading edge of protrusions, predominantly beneath the lamellipodium Choi et al. [2008], Wolfenson et al. [2009], and are linked to a filamentous actin meshwork Lock et al. [2008].

Nascent adhesions are believed to be responsible for generating and transmitting significant traction forces that propel migrating fibroblasts forward Beningo et al. [2001]. Small nascent adhesions are also associated with the rapid migration of other cell types, such as CHO.K1 cells Choi et al. [2008], and with the control of cell migration through FAK-talin interactions Lawson et al. [2012]. While studied more extensively in 2D migration, nascent adhesions have also been shown to play a crucial role in migration within 3D environments. Highly dynamic nascent adhesions, exhibiting rapid assembly and disassembly with an average lifetime of approximately one minute, have been observed in U2OS osteosarcoma cells migration in 3D matrices Kubow and Horwitz [2011]. The importance of nascent adhesions in both 2D and 3D migration underscores the need to understand their regulation.

Rigorous experimental investigations, complemented by mathematical and computational studies, have aimed to elucidate the dynamics of adhesions. Nevertheless, integrin-based adhesions to the ECM are highly complex processes involving multiple stages, each characterized by distinct molecules and functions that contribute to specific mechanisms.

To fully understand adhesion mechanisms, it is necessary to dissect each stage and treat it uniquely due to the different components and dynamics involved. For nascent adhesions, the earliest adhesion structures to appear during cell migration, while it is acknowledged that the mechanisms of their assembly (nucleation) and disassembly are not yet fully understood Parsons et al. [2010], it has been observed that actin motor proteins are absent in these structures Choi et al. [2008]. Actin motor proteins such as myosin II are essential for generating tension along actin fibers, which promotes the growth and maturation of small adhesions into larger focal adhesions. Myosin II has been observed in focal complexes, structures larger than nascent adhesions. Nascent adhesions have also been reported to nucleate on soft substrates, *i*.*e*., ECM with compliant stiffness, whereas more rigid substrates trigger the growth and maturation of adhesions Mason et al. [2012].

Existing mathematical models have primarily focused on the interactions between nascent adhesion dynamics and intracellular components such as actin cytoskeleton dynamics and barbed ends Choi et al. [2008], actin and myosin dynamics Welf et al. [2013], adhesion and membrane protrusion Cirit et al. [2010], and actin organization and adhesion Shemesh et al. [2012]. While valuable for studying adhesion, these models often lack the inclusion of ECM dynamics, whose rigidity regulates adhesion formation and maturation. Some models do integrate the mechanics (rigidity) of the ECM and intracellular dynamics Pathak and Kumar [2011], Walcott et al. [2011] but these models are often too general to specifically address nascent adhesions, particularly given the absence of myosin II involvement. This raises the question of whether the formation of specific nascent adhesion profiles can be explained by a bi-directional pathway specific to nascent adhesions, incorporating the dynamics of both intracellular and extracellular environments. Another question is what kind of influence the local extracellular environment exerts on the assembly and/or disassembly of nascent adhesions. In this paper we present a mathematical model for the dynamics of nascent adhesion resulting from its interactions with the actin cytoskeleton and the ECM. These three components are modeled as interconnected compartments with a time delay from ECM to adhesion, represented by a system of ordinary differential equations with time delay. The model solutions demonstrate that the temporal profiles of nascent adhesions obtained from experimental studies by Choi et al in 2008 Choi et al. [2008] can be predicted without invoking spatial dynamics, solely by considering the concentration changes of the three components over time. Our computational results qualitatively match the experimental data.

## 2 The Model

As the earliest identifiable adhesion structures, nascent adhesions are considered crucial in cell migration, being present in various cell types during the initial stages of motility on 2D substrates and in some cells within 3D environments. Understanding the mechanisms governing nascent adhesion assembly (or nucleation) and disassembly (or turnover), which ultimately dictate the nascent adhesion profile, is essential for elucidating the multi-step processes of cell migration. In this section we describe our mathematical model, informed by experimental studies on early-stage migration. Our model considers three key components as our mathematical variables: nascent adhesions, the intracellular cytoplasm (specifically focusing on actin dynamics), and the ECM. We hypothesize that these components interact as a system of interconnected compartments through which adhesion signaling flows, as depicted in Fig. 2.

**Figure 2:**
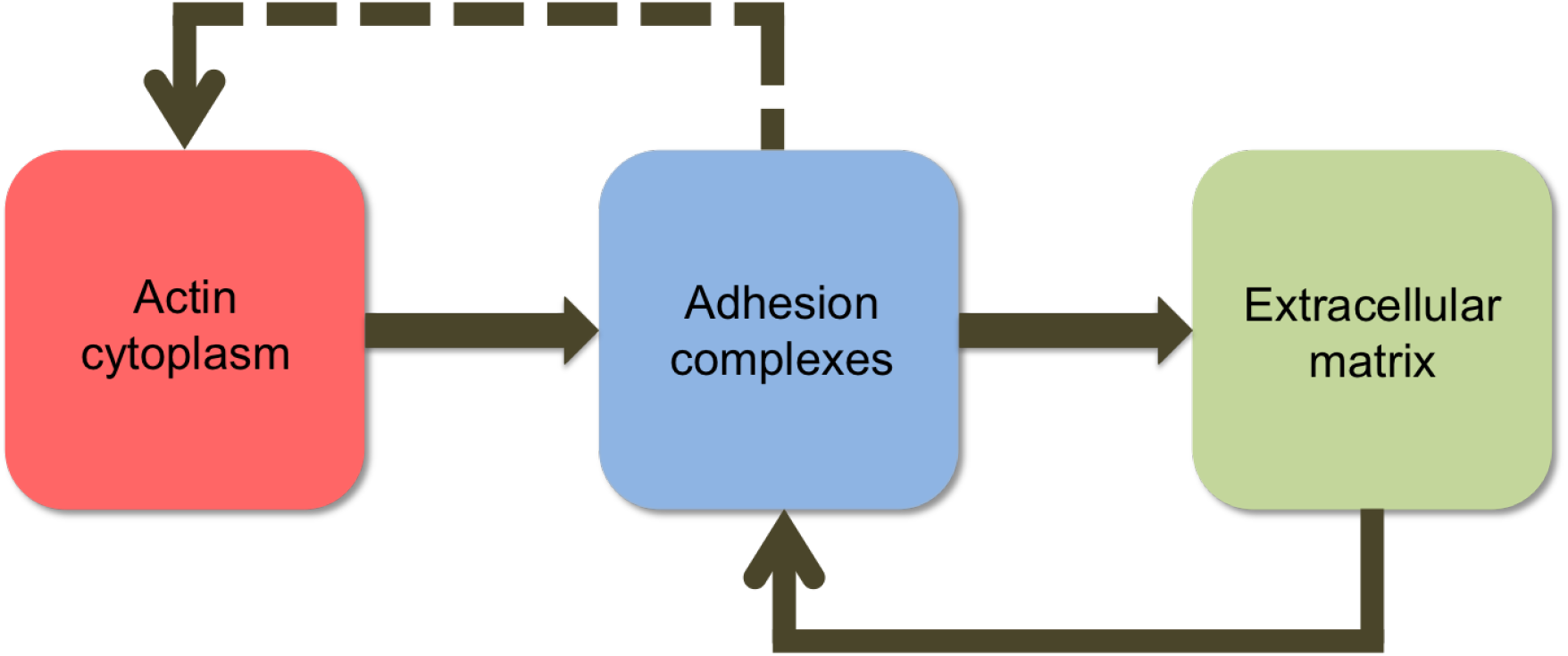
Schematic diagram illustrating the compartmental model describing the interactions between adhesion, cytoskeleton dynamics, extracellular matrix stiffness to produce adhesion signaling required for cell migration.

In Fig. 2, we postulate that nascent adhesion nucleation is driven by intracellular compartment dynamics, specifically the actin cytoskeleton Parsons et al. [2010]. The formed adhesions then exert force on the nearby ECM through ligand binding with integrin clusters (not explicitly modeled here but assumed to be included in the ECM compartment). Adhesion formation is sensitive to ECM stiffness, manifesting as a feedback mechanism from the ECM to adhesion sites, which is subsequently transduced into intracellular biochemical signals. ECM stiffness influences the size of adhesions, contributing to nascent adhesion nucleation. Once adhesions reach a sufficient size, involving a greater number of protein components, they exert both mechanical and chemical effects on actin filaments within the cytoskeleton Gardel et al. [2010], Wolfenson et al. [2009] (represented by a dashed line in Fig. 2), an aspect we do not explicitly model in this paper.

### 2.1 Actin Cytoskeleton

The actin cytoskeleton is a major component of the cell cytoplasm and plays critical roles in numerous intracellular processes. Simplifying the intricate details of actin dynamics, within the cytoplasm compartment, we focus solely on actin reorganization. This process is rapid, involving the polymerization of actin monomers into filaments and the subsequent depolymerization back into monomers. Polymerization, leading to actin filament growth, occurs predominantly at the plus (or barbed) end, while depolymerization, resulting in filament disassembly, occurs at the minus (or pointed) end.

Beyond assembly/polymerization and disassembly/depolymerization, the dynamics of actin filaments are also influenced by retrograde flow Choi et al. [2008], Alexandrova et al. [2008]. For simplification, we assume that actin organization dynamics consist solely of assembly/polymerization and disassembly/depolymerization. Following Dawes et al. [2006], Dawes and Edelstein-Keshet [2007], we model actin dynamics using first-order kinetics for filament growth due to monomer addition at barbed ends with a rate *k*_1_. For this purpose, we define the state of the cytoskeleton compartment by the density of actin filament length, denoted by *a*(*t*), and the density of barbed ends, denoted by *b*(*t*). To account for the rapid actin filament dynamics observed *in vivo* Ono [2007], *we model disassembly with a second-order kinetic term with a rate k*_2_. To prevent excessive disassembly and ensure filament formation, we also assume that monomer addition at barbed ends concurrently limits rapid disassembly with the same order of kinetics. Consequently, we model the concentration of actin filament length *a*(*t*) as:

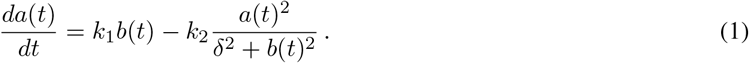

where *δ*is a small positive constant (0 *≤ δ≤* 1) introduced to account for more effects of molecule acceleration. The disassembly term can be interpreted as containing a spatial-independent retrograde flow, implicated as a contributing factor to actin filament depolymerization or disassembly. The dynamics of barbed ends *b*(*t*) are given by:

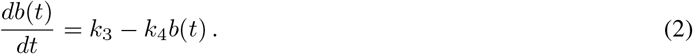

Equation 2 is a simplified representation of more complex barbed end dynamics, where new barbed ends are initiated by Arp2/3 Mogilner and Edelstein-Keshet [2002]. In our model, the rate of barbed end initiation is represented by a constant *k*_3_. Barbed ends are diminished due to capping by capping proteins, modeled here as a first-order degradation process with a rate *k*_4_. Equation 2 is an inhomogeneous first-order ordinary differential equation which can be solved analytically on its own:

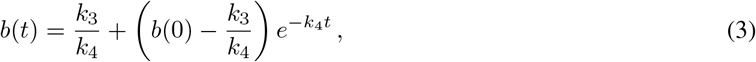

where *b*(0) is the barbed end concentration at the initial time *t* = 0. If *k*_3_*/k*_4_ *< b*(0), the solution is a decreasing function, contributing to the fast disassembly of actin filaments.

### 2.2. Extracellular Matrix

The ECM is a crucial environment that provides cells with chemical and mechanical stimuli that influence cell behavior and fate. The binding of integrins to ECM ligands enables cells to exert tension on the surrounding ECM Ilić et al. [2004]. For large, mature adhesions like focal and fibrillar adhesions, their size has been shown to correlate linearly with cell-generated forces, where the size increases and decreases as a function of applied force Mason et al. [2012]. However, smaller adhesions exhibit a different relationship with force, tending to match the forces they exert on the ECM to the ECM’s stiffness. Nascent adhesions and focal complexes can exert appreciable forces on the ECM, ranging from picoNewtons (pN) to nanoNewtons (nN) Bruinsma [2005]. Conversely, the ECM’s Young’s modulus represents the rigidity or stiffness sensed by nascent adhesion sites. Nascent adhesions (*<* 1 *µm*^2^) typically form on soft (compliant) ECM (*∼* 1 kPa), while larger adhesions like focal adhesions are found on substrates with higher stiffness (*∼* 30 – 100 kPa) Mason et al. [2012], Sarvestani [2011].

The binding of integrins with ECM ligands or macromolecules on the cell surface imposes spatial restrictions on signaling and ECM dynamics Humphries et al. [2006]. Forces exert on the ECM by adhesions cause displacement of ECM components. At the smaller scale of nascent adhesions with picoNewton-range, forces, these forces primarily affect the displacement at integrin-ligand binding sites, representing the closest interaction between adhesion sites and the ECM. According to elasticity theory, this displacement is inversely proportional to the ECM’s Young’s modulus. In this paper, we model this displacement as the key component of the ECM compartment. Similar to our treatment of actin concentration, our model simplifies detailed molecular-level processes within the ECM compartment. The ECM displacement is assumed to be governed by the binding of integrins in the adhesion compartment with ECM ligands, formulated as a nonlinear delay differential equation:

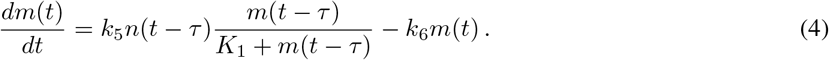

Here, *m*(*t*) represents the ECM displacement, *k*_5_ is the rate of displacement arrangement due to adhesion formation, and *K*_1_ is a saturation constant that limits the displacement (estimated as *K*_1_ = 1). The displacement is assumed to decay with a rate *k*_6_. The parameter *τ* represents a time delay, which can be interpreted as the mechanical relaxation time if we consider integrin-ECM bonds as viscoelastic material, reflecting the time lag in ECM displacement in response to adhesion forces.

### 2.3 Adhesion

Despite extensive research over the past three decades, the precise mechanisms underlying nascent adhesion nucleation, elongation, and disassembly remain incompletely understood. Nevertheless, two primary models have been proposed for nascent adhesion nucleation Parsons et al. [2010]: (i) nucleation or assembly initiated by the binding of integrins to ECM proteins, clustering of integrin-ligand complexes, and subsequent recruitment of new adhesion components to their clustered cytoplasmic domains; and (ii) nucleation or assembly initiated by actin polymerization at dendritic actin networks. Here, we attempt to incorporate elements of both models. Consistent with the second model and experimental observations, we assume that the assembly of nascent adhesions in the lamellipodium requires actin polymerization Choi et al. [2008]. Adhesions of all sizes serve as anchors for actin filaments undergoing retrograde flow towards the cell membrane. We model this assembly process using first-order kinetics with a rate *k*_7_, proportional to the actin filament length density *a*(*t*).

In addition to being influenced by intracellular components, nascent adhesion assembly is also regulated by the extracellular matrix. For the second term, we assume that nascent adhesion growth is limited by the extent of molecular ECM displacement. We model a non-linear adhesion growth term with a first-order kinetic rate *k*_8_, which is negatively regulated by the ECM displacement *m*(*t*) through a saturation term. The rate of change of nascent adhesion density *n*(*t*) is therefore formulated as:

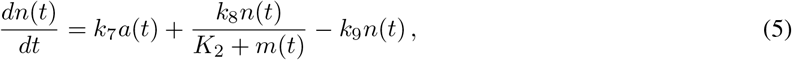

where adhesion disassembly is modeled as a first-order removal process with a rate *k*_9_, and *K*_2_ is a saturation constant (set to *K*_2_ = 1).

## 3 Parameter Estimation

To estimate the parameter values for our mathematical model using experimental data obtained from the literature, we employed the `ParameterOptimizationToolkit` in MATLAB®. This toolbox provides a collection of heuristic algorithms designed to find a set of parameters that yields model outputs closely matching observed data.

The optimization toolbox offers several algorithms suitable for refining initial parameter guesses, including gradient descent, iterated local search, simulated annealing, and genetic algorithms. Among these, simulated annealing provided the best approximation for fitting our model to the experimental data. Simulated annealing effectively avoids entrapment in local optima by employing an acceptance probability criterion for energy minimization, analogous to the Metropolis algorithm:

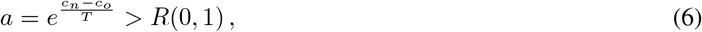

where *a* is the acceptance probability, (*c*_*n*_ − *c*_*o*_) represents the difference between the new cost *c*_*n*_ and the old cost *c*_*o*_, *T* is the system temperature, *e* is the base of the natural algorithm (*≈* 2.71828), and *R*(0, 1) is a random number drawn from a uniform distribution in the interval [0, 1]. A key feature of this algorithm is its ability to accept not only new parameter sets that lower the objective function (cost), but also, with a certain probability, those that increase it. This allows the algorithm to escape local optima during early iterations and explore the parameter space more globally for better solutions. Using this algorithm, we obtained the parameter values listed in Table 1.

**Table 1:** Estimated parameter values.

| Parameter | Value used in model | Experimental range | Model dimension | Reference |
| --- | --- | --- | --- | --- |
| $k_1$ | 0.15 | $0.5 \mu\text{m s}^{-1}$ | $\text{s}^{-1}$ | Dawes and Edelstein-Keshet [2007] |
| $k_2$ | 0.023 | 0.0083 | $\mu\text{m} \mu\text{m}^{-1} \text{s}^{-1}$ | Choi et al. [2008] |
| $k_3$ | 0.091 | not known | $\mu\text{m}^{-1} \text{s}^{-1}$ | estimated |
| $k_4$ | 0.01 | 1 | $\text{s}^{-1}$ | Mogilner and Edelstein-Keshet [2002] |
| $k_5$ | 0.5 | not known | $\text{M s}^{-1}$ | estimated |
| $k_6$ | 0.02 | not known | $\text{M s}^{-1}$ | estimated |
| $k_7$ | 0.05 | $0.05 - 0.0667$ | $\text{s}^{-1}$ | Choi et al. [2008] |
| $k_8$ | 0.75 | not known | $\text{s}^{-1}$ | estimated |
| $k_9$ | 0.035 | $0.000167 - 0.0667$ | $\text{s}^{-1}$ | Choi et al. [2008], Cirit et al. [2010] |
| $\tau$ | 95.0 | $> 20$ | s | Crow et al. [2012] |

The experimental data for adhesion and actin densities were extracted from the work of Choi et al. Choi et al. [2008], who observed nascent adhesions near the leading edge of protrusions in migrating cells. These small adhesions consistently form at the periphery of active protrusions, undergoing rapid assembly and disassembly with an average lifespan of 76.1 ± 22 s and a transient phase duration of 11.8 ± 6.2 s Choi et al. [2008]. Their experiments also demonstrated that the rate of nascent adhesion formation is directly coupled to the rate of protrusion. The specific data extracted from Choi et al. Choi et al. [2008] for parameter fitting of our model are presented in the Results section.

## 4 Parameter Sensitivity Analysis

In this section, we perform a local sensitivity analysis to assess the robustness of our model outputs to small perturbations in the input parameters around the default parameter values listed in Table 1.

We can express the system of equaitons 13 in a compact vector form as:

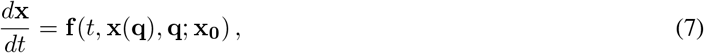

where *t* represents time, **x**(**q**) *∈* ℝ^*n*^ is the vector of *n* system outputs (state variables or solutions), **q** *∈* ℝ^*m*^ is the vector of *m* parameters, and **x**_**0**_ is the vector of initial conditions. The sensitivity functions are defined as the partial derivatives of the state variables with respect to the parameters. Differentiating Equation 7 with respect to **q** yields the following sensitivity differential equations Turanyi et al. [1988]:

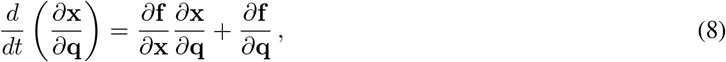

where 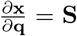 are the local sensitivity coefficients. In our case, these correspond to the sensitivity of actin filament length (*A*), ECM displacement (*M*), and nascent adhesion density (*N*) with respect to the parameter vector **q**:

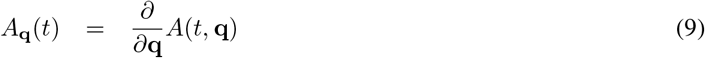

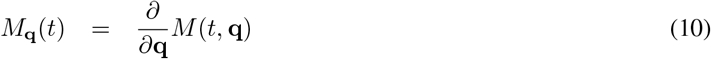

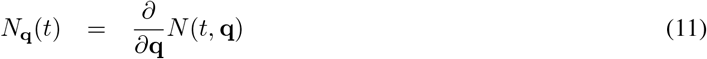

after replacing differential notation with subscript notation, *i*.*e*.,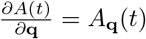.

To obtain numerical approximations of these local sensitivity coefficients Eq. 9-11, each parameter value was perturbed by ±1%. Following the approach outlined in Nagaraja et al. [2014], the derivative was approximated using the second-order central finite difference method and we adjusted the MATLAB’s sensitivity toolbox provided in their supplementary information accordingly. The local sensitivity coefficients were then calculated by the relative local sensitivity formula given by

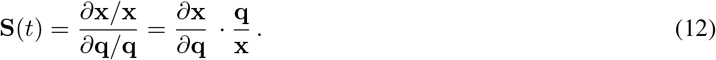

The results of this analysis are presented in Tables 2 and 3 for the parameter sets **q** = [*k*_1_, *k*_2_, *k*_3_, *k*_4_, *k*_5_, *k*_6_, *k*_7_, *k*_8_, *k*_9_]^*T*^ and **q** = [*δ, K*_1_, *K*_2_]^*T*^, respectively, both for the time span *t* = 1000 s. For nascent adhesion density, the parameters *k*_1_, *k*_2_, *k*_3_, *k*_4_, and *k*_7_ exhibit the highest sensitivity, showing immediate effects upon perturbation due to their direct involvement in actin polymerization and initial adhesion formation. The parameter *k*_9_ (adhesion disassembly rate) shows intermediate sensitivity. Perturbing *δ*results in an insignificant effect, likely due to the small magnitude of its variation (10^*−*3^ order). The remaining parameters (*k*_5_, *k*_6_, *k*_8_, *K*_1_, *K*_2_ and *τ*) demonstrate lower sensitivity, exhibiting an instantaneous response to perturbation followed by a rapid decay towards zero. The ECM displacement shows similar sensitive parameters (*k*_1_, *k*_2_, *k*_3_, *k*_4_, and *k*_7_) and an expanded set of semi-sensitive parameters (*k*_6_, *k*_9_, and *K*_1_). The remaining parameters (*k*_5_, *k*_8_, *K*_2_, and *τ*) have a less pronounced impact on ECM displacement. Actin filament length density is primarily affected by *k*_1_, *k*_2_, *k*_3_, and *k*_4_, all of which show significant sensitivity. The parameters governing barbed end dynamics (*k*_3_ and *k*_4_) were found to be robust to perturbation and are therefore not explicitly shown in the sensitivity analysis figures.

**Table 2:**
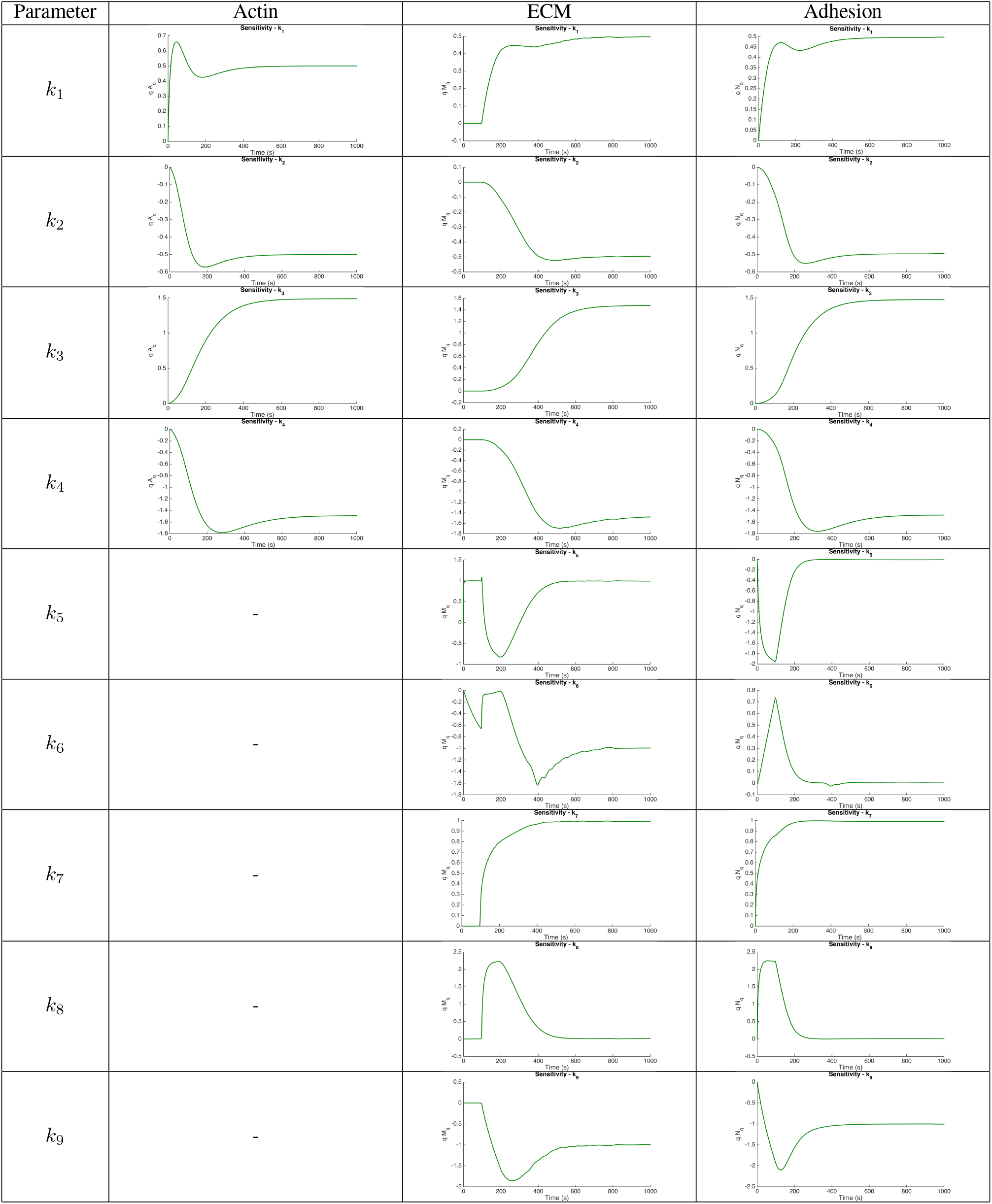
Results of parameter sensitivity 1.

**Table 3:**
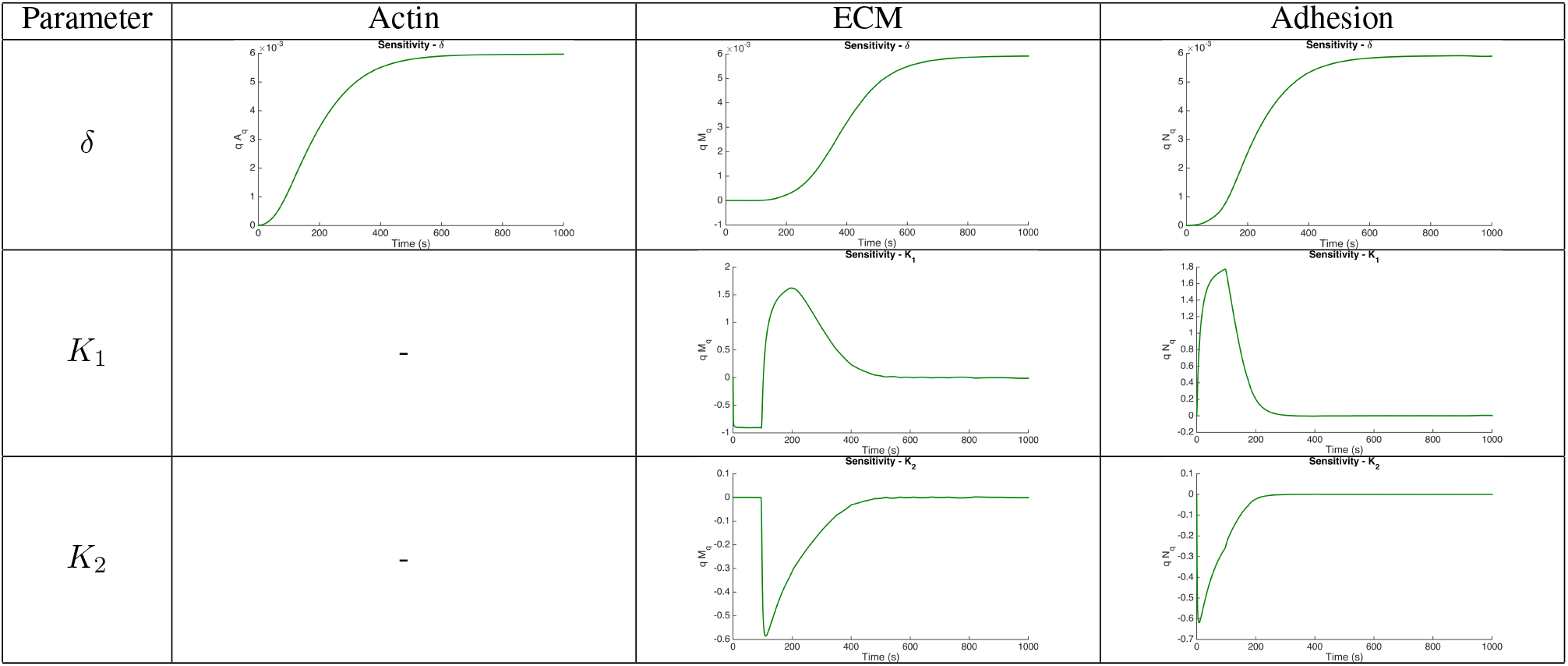
Results of parameter sensitivity 2.

| Parameter | Actin | ECM | Adhesion |
| --- | --- | --- | --- |
| $\delta$ | | | |
| $K_1$ | - | | |
| $K_2$ | - | | |

## 5 Model Analyses

### 5.1 Local Stability Analysis

In this analysis, we seek to determine the existence and stability of equilibrium points for our model equations, which are rewritten as:

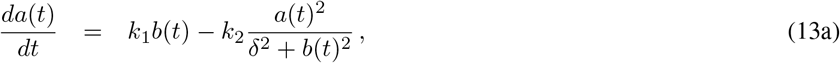

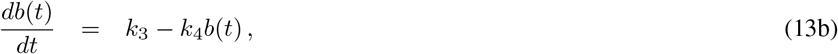

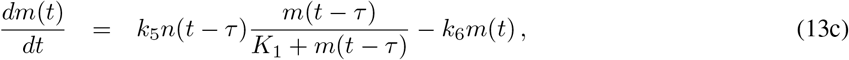

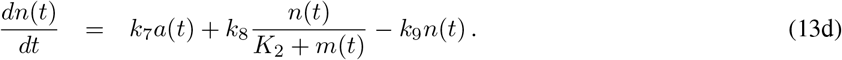

Assuming *δ*= *K*_1_ = *K*_2_ = 1 for simplicity in finding steady states, the system of equations 13 yields potential steady state points, **u**_ss_ = (*a*_ss_, *b*_ss_, *m*_ss_, *n*_ss_), determined by setting the time derivatives to zero:

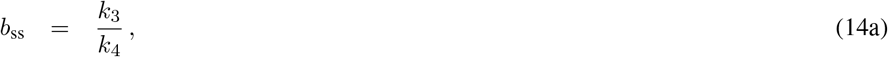

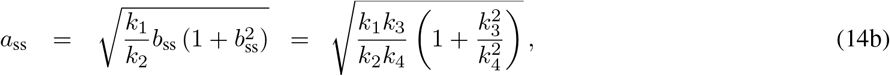

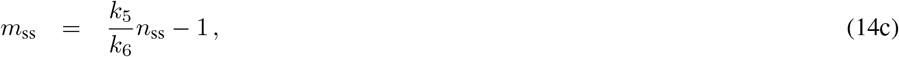

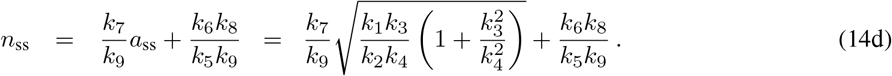

Using the parameter values provided in Table 1, we find a single positive non-zero steady state point: **u**_*ss*_ = (70.526, 9.1, 2539.2, 101.609). To investigate the stability of this steady state, we introduce small perturbations around it: *ã*, 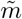, and *ñ*, such that the system solutions are given by:

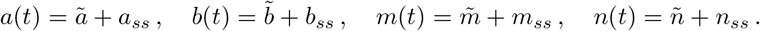

Linearizing the system of equations 13 around this steady state yields:

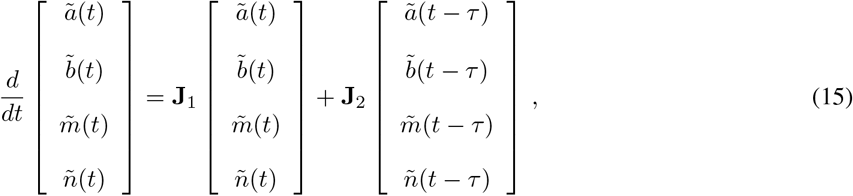

where **J**_1_ and **J**_2_ are 4 × 4 Jacobian matrices evaluated at the steady state:

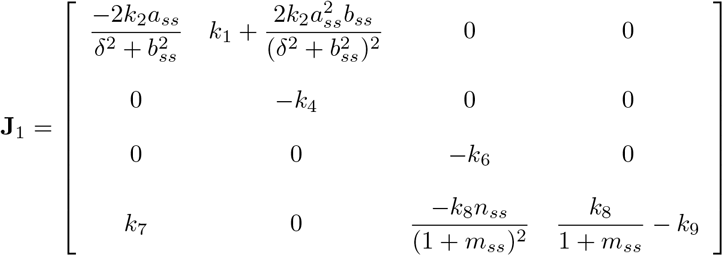

and

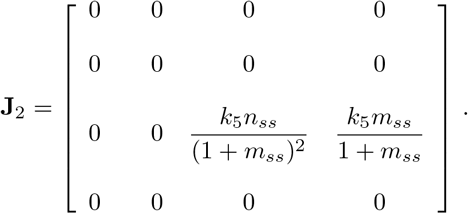

The stability of the steady state is determined by the eigenvalues *λ* of the characteristic equation:

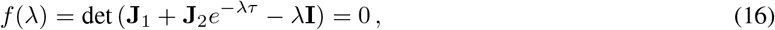

which results in the following fourth-order polynomial:

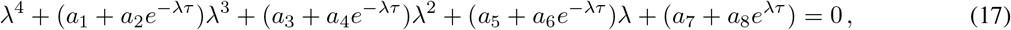

where the coefficients *a*_*i*_ are defined as:

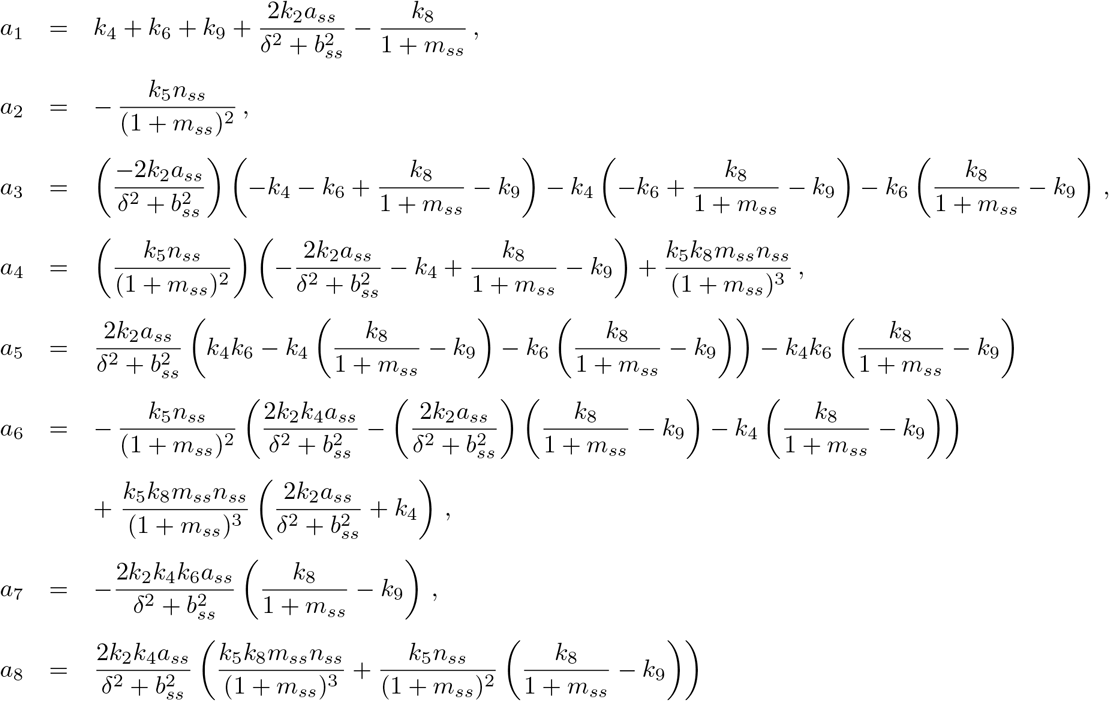

If *τ >* 0, the system of equations becomes difficult to solve because of the following reasons: (i) with non-zero delay *τ >* 0, we now deal with a transcendental equation that has trigonometric functions which implies that it has infinitely many eigenvalues and the classical Routh-Hurwitz conditions therefore cannot be applied, and (ii) some general tests to determine when all eigenvalues of the transcendental equations have negative real parts are very complicated to apply and far from trivial.

However, based on the transcendental analysis in Murray [2003], the real parts ℝ (*λ*) of all solutions of Eq. 17 are bounded above. To analyze the stability, we investigate the existence of purely imaginary roots Li et al. [1999], Culshaw and Ruan [2000], as a change in stability occurs when an eigenvalue crosses the imaginary axis.

We set *λ*(*τ*) = *µ*(*τ*) + *iν*(*τ*) with *ν >* 0, where *µ*(*τ*) and *ν*(*τ*) are delay-dependent of the real and imaginary part, respectively. By continuity, if the time delay *τ >* 0 is sufficiently small, the real part *µ*(*τ*) is negative and the steady state **u**_ss_ is stable. When *µ*(*τ*) transitions to positive, there is a critical delay *τ*_0_ *>* 0 at which *µ*(*τ*_0_) = 0 that makes *λ* = *iν*(*τ*_0_) a purely imaginary root of Eq. 17, then the steady state loses its stability and becomes unstable when *µ*(*τ*) *>* 0. If such *ν*(*τ*_0_) does not exist or if Eq. 17 does not have purely imaginary roots for all delay, the steady state is always stable. Below we shall show that this is true for Eq. 17.

Substituting *λ* in Eq. 17 with *µ*(*τ*) + *iν*(*τ*), the characteristic equation becomes:

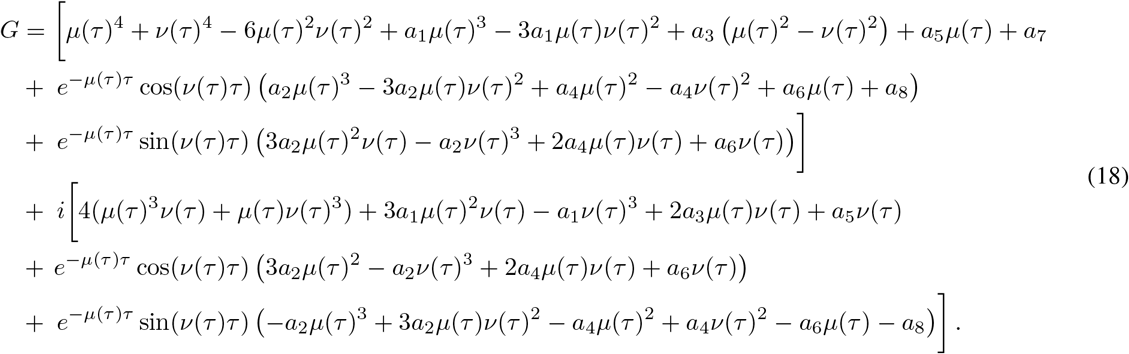

For purely imaginary roots, by setting *µ*(*τ*) = 0, we have *iν*(*τ*) a root of *G* = 0. Separating the real and imaginary parts in Eq. 18 yields:

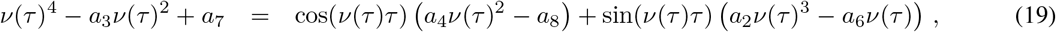

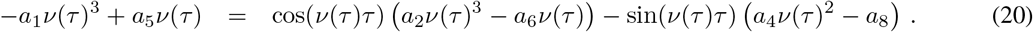

If temporarily we let *A* = *a*_4_*ν*(*τ*)^2^ − *a*_8_ and *B* = *a*_2_*ν*(*τ*)^3^ − *a*_6_*ν*(*τ*), taking square of both sides gives:

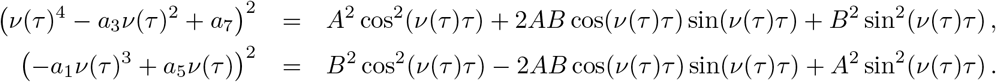

Adding the two terms up we get:

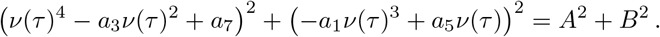

Now, substituting *A* and *B* back and expanding the left hand side leads to the 8-th order polynomial:

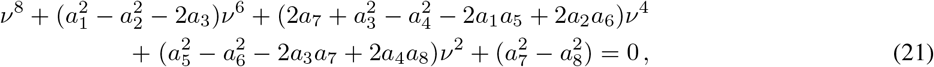

after replacing *ν*(*τ*) with *ν* for simplification.

If we let *ω* = *ν*^2^, the order of polynomial reduces four degrees, where now Eq. 21 becomes:

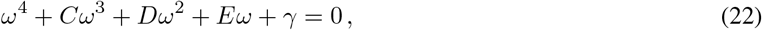

and

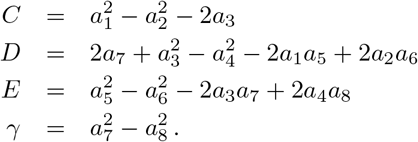

Using the parameter values from Table 1, the coefficients of Eq. 22 are approximately *C* = 0.0032, *D* = 4.576 × 10^*−*6^, *E* = 3.2 × 10^*−*10^, and *γ* = 7.22 × 10^*−*14^, where *D, E*, and *γ* are very small, and hence, can be neglected.

If we consider *D* = 0, *E* = 0, and *γ* = 0, the Eq. 22 becomes:

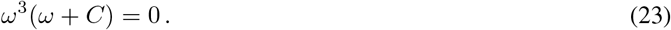

The roots of Eq. 23 are *ω* = 0 and *ω* = − *C* = − 0.0032. It follows that Eq. 21 has no positive roots. This implies that there is no *ν* such that *iν* is an eigenvalue of the characteristic equation Eq. 17. Therefore, the real parts of all the eigenvalues of Eq. 17 are zero for all time delay *τ* ≥ 0, and the steady state **u**_ss_ is stable.

### 5.2 Global Stability Analysis

Let a linear system of delay differential equations given by

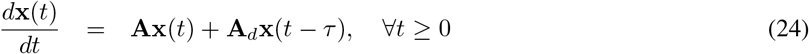

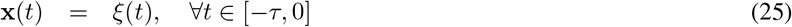

where **x**(*t*) *∈* ℝ^*n*^ is a system state vector, *ξ*(*t*) is a system initial condition function, **A, A**_*d*_ *∈* ℝ^*n×n*^ are constant matrices (transition matrices), and *τ* : 0 *≤ τ ≤ τ*_*M*_ is a discrete delay upper bounded by *d*_*M*_ .

Let the state vector **x**(*t*) = [*a*(*t*) *b*(*t*) *m*(*t*) *n*(*t*)], hence, the linearized transition matrices are given by

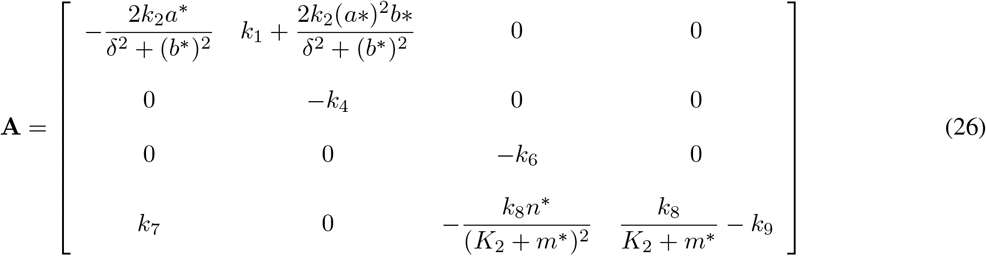

and

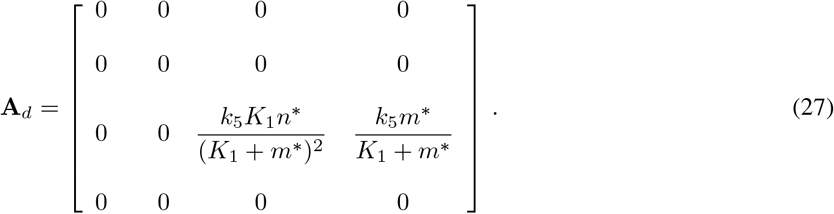

The following theorem guarantees the delay dependent stability for the system in Eq. 24.

#### 5.2.1 Discrete Delay Stability Analysis

##### Theorem 1

*Gouaisbaut and Peaucelle [2006] A time delay system (24) has asymptotic stability for any delay inside* 0 ***≤*** *τ* ***≤*** *τ*_*M*_ *if there exist matrices* **P *∈*** ℝ^*n×n*^ *>* 0, **Q *∈*** ℝ^*n×n*^ *>* 0 *and* **R *∈*** ℝ^*n×n*^ *>* 0, *such that the following linear matrix inequality (LMI) holds*

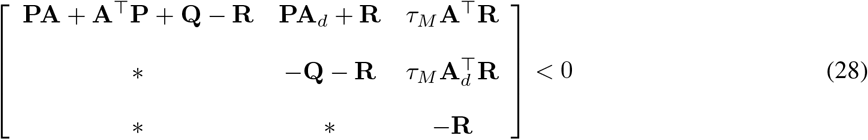

**Proof 1** *The following Lyapunov-Krasovskii functional (LKF) for stability analysis of system Eq. 24 is used*

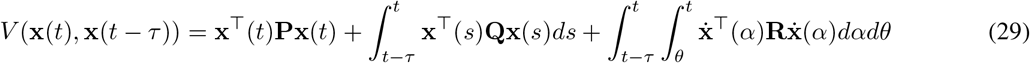

*Taking the derivative of LKF along trajectories of the system Eq. 24 gives*

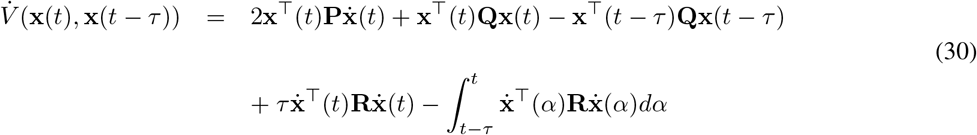

*The integral term can be bounded by using Jensen’s inequality. The reader may refer to Gouaisbaut and Peaucelle [2006] for this proof*.

The nascent-adhesion system has an equilibrium point at *a*^*\**^ = 70.526, *b*^*\**^ = 9.1, *m*^*\**^ = 2539.2, and *n*^*\**^ = 101.609. Applying Gouaisbaut’s theorem Gouaisbaut and Peaucelle [2006] to the linearized nascent-adhesion system, we obtain the following condition matrices

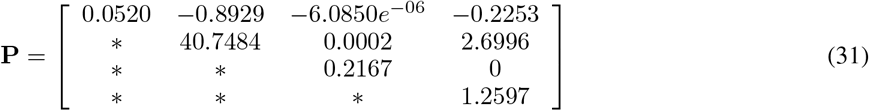

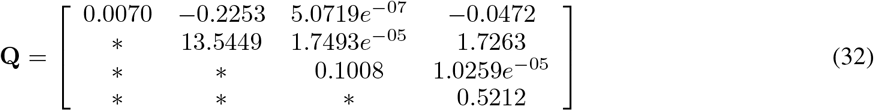

and

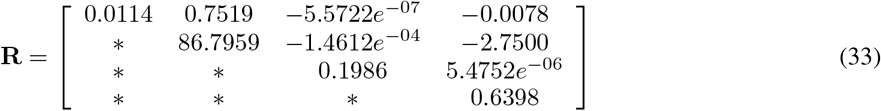

for *τ*_*M*_ = 20 (see the Numerical Simulations section on how this value was obtained). The matrices **P, Q**, and **R** are positive definite, therefore, it can be concluded that the nascent-adhesion system is stable at the equilibrium point.

#### 5.2.2 Continuous Time Varying Delay Stability Analysis

A nascent-adhesion system may experiencing in a varying time delay such as *τ* (*t*). Therefore, the system can be presented as

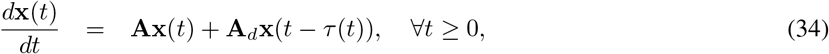

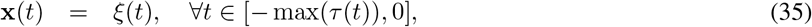

for *τ* (*t*) : 0 *≤ τ* (*t*) *≤ τ*_*M*_ .

##### Theorem 2

*Jing et al. [2004]* **A delayed system presented in equation (34) is asymptotically stable if there exists matrices P** *>* 0, **Q** *>* 0, **R, S** *and* **X**_*ij*_, *i ≤ j, i, j* = 1, 2, 3 *such the following conditions are satisfied*

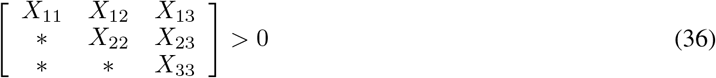

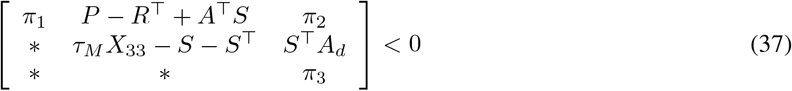

*where π*_1_ = 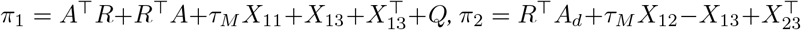 and 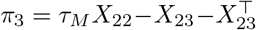

**Proof 2** *Let consider a LKF as the following*

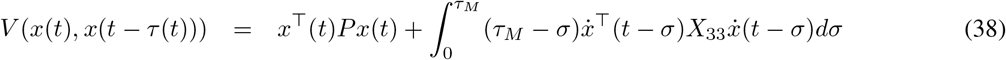

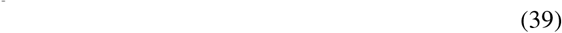

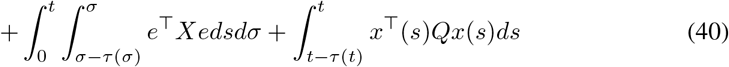

*where*

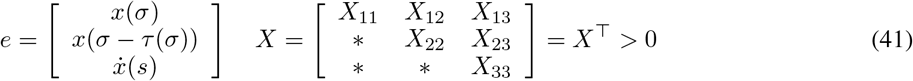

*by considering that (34) becomes*

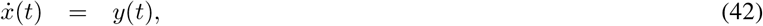

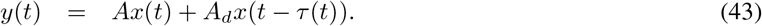

*Having derivative along the trajectories of a system (34) for with the derivative of this term has*

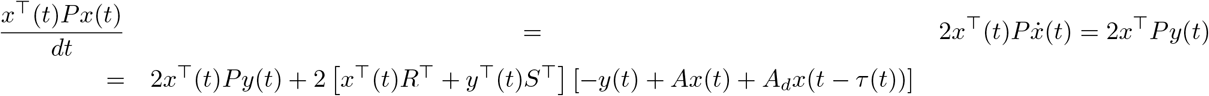

*derivative other terms can refers to Jing et al. [2004] for complete proof*.

After applying Jing’s theorem 2, it is found that

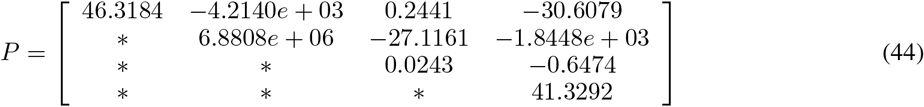

and

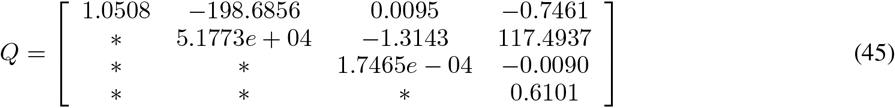

are positive definite. Therefore, the nascent-adhesion system is asymptotically stable at the linearized point.

## 6 Numerical Simulations

In this section we solve the system of differential equations 1, 2, and 5 along with the delay differential equation 4 employing the parameter values listed in Table 1. The following initial conditions were assumed for actin filament density (*a*), barbed end density (*b*), ECM displacement (*m*), and adhesion density (*n*):

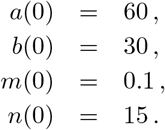

These initial conditions represent a scenario where a baseline level of actin filaments is present at the site of nascent adhesion formation (or at the leading edge of the lamellipodium), while the initial adhesion concentration, represented by unbound or talin-bound integrin clusters, is relatively low. We assume that significant ECM displacement has not yet occurred at the onset of adhesion assembly, hence its initial value is set to a small value (*<* 1).

The numerical solutions presented below illustrate the temporal dynamics of *a*(*t*), *m*(*t*), and *n*(*t*) as they approach the stable steady state **u**_ss_ = (*a*^*\**^, *b*^*\**^, *m*^*\**^, *n*^*\**^), which was confirmed through stability analysis presented in the previous section. To facilitate comparison with the normalized experimental data from Choi et al. Choi et al. [2008], the model solutions were normalized by their respective maximum values.

Figure 3 displays the temporal evolution of nascent adhesion density *n*(*t*). The solid blue line represents the model solution, while the blue dots correspond to the experimental data. Adhesion assembly commences at *t* = 0 and continues until *t* = 95 s, which corresponds to the time delay *τ* in Eq. 4 and also aligns with the observed maximum assembly time of nascent adhesions before the onset of disassembly/turnover in the experimental data. The dashed gray line in Fig. 3 illustrates the model solution for adhesion when the second term on the right-hand side of Eq. 5 is omitted, effectively removing the influence of ECM displacement on adhesion dynamics. This comparison highlights that the initial growth of nascent adhesions is primarily driven by actin filament polymerization. As adhesions mature, forces at the adhesion site accumulate. After the time delay *τ* = 95 s, the effects of displacement, resulting from these adhesion-generated forces, become significant, leading to the initiation and acceleration of nascent adhesion disassembly. Increased displacement, associated with softer ECM, further enhances adhesion turnover. These model predictions are consistent with experimental findings indicating that nascent adhesions grow and turn over more rapidly on softer substrates. Furthermore, the overall profile of nascent adhesion predicted by our model, encompassing assembly, transient stability, and disassembly phases, qualitatively matches the literature data. The duration of the transient stability phase in our model is on the same order as that reported in Choi et al. [2008] (11.8 ± 6.2 s).

**Figure 3:**
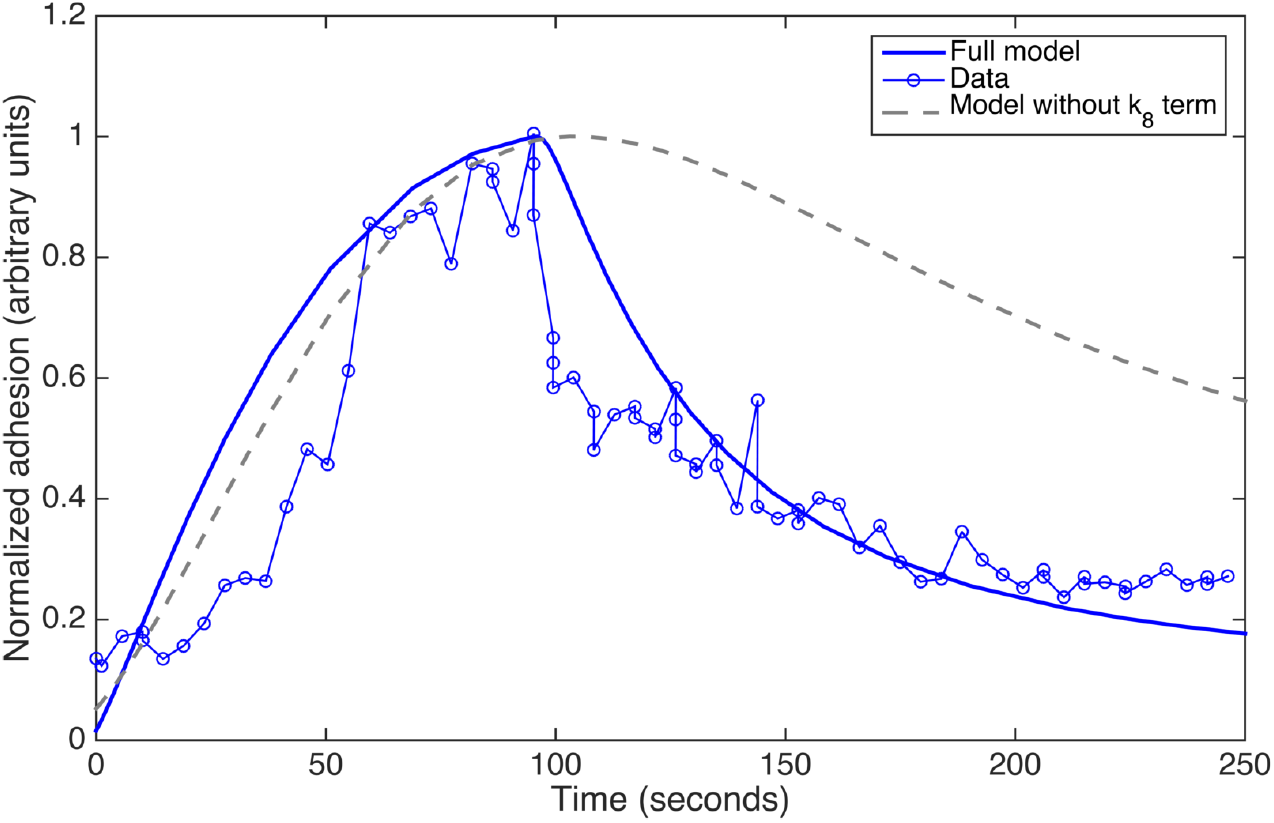
Plots showing the comparison between computational result of our model (solid line) and data (dots) from Choi et al. [2008] for a nascent adhesion profile in arbitrary units, both were simulated and sampled until 250 s. The model solution is in agreement and follows the experimental data trend. Dashed line represents the solution of model Eq. 5 without the effect of ECM stiffness on adhesion dynamics. Data is adapted by permission from Macmillan Publishers Ltd: Nature Cell Biology. Choi et al, Nature Cell Biology, 10, 9 (2008), copyright 2008.

Figure 4 presents the temporal dynamics of actin filament concentration *a*(*t*) (solid line) simulated over 250 seconds, along with the corresponding experimental data (dots). The actin filament dynamics exhibit a similar trend to adhesion, characterized by an initial increase in concentration, followed by a slight stabilization and subsequent decrease. Although the model solution appears slightly broader or shifted compared to the data points, the overall trend is remarkably similar. This dynamic behavior arises from the simplified kinetics of actin polymerization, influenced by barbed end dynamics, which are also modeled with basic kinetic terms. While our model represents a reduced version of the complex spatial and regulatory mechanisms governing actin polymerization in cells, it appears sufficient to capture the essential temporal dynamics relevant to nascent adhesion assembly. Studies suggest a direct link between the Arp2/3 complex in the lamellipodial actin network and nascent adhesion components (reviewed in Wolfenson et al. [2009]). In our model, this connection is indirectly represented in Eq. 2, where a constant rate *k*_3_ models the initiation of new barbed ends (a process often mediated by Arp2/3), and subsequent actin polymerization occurs through monomer addition at these barbed ends. This simplification is adequate to support the linear kinetic term in our adhesion model (the first term on the right-hand side of Eq. 5), which posits that adhesion assembly is connected to and driven by actin polymerization. Furthermore, the transient dynamics of actin filaments during nascent adhesion formation predicted by our model are consistent with hypotheses proposed in a review by Ciobanasu et al. Ciobanasu et al. [2012] regarding transient interactions between *α*-actinin and integrin *β*_1_ within nascent adhesions.

**Figure 4:**
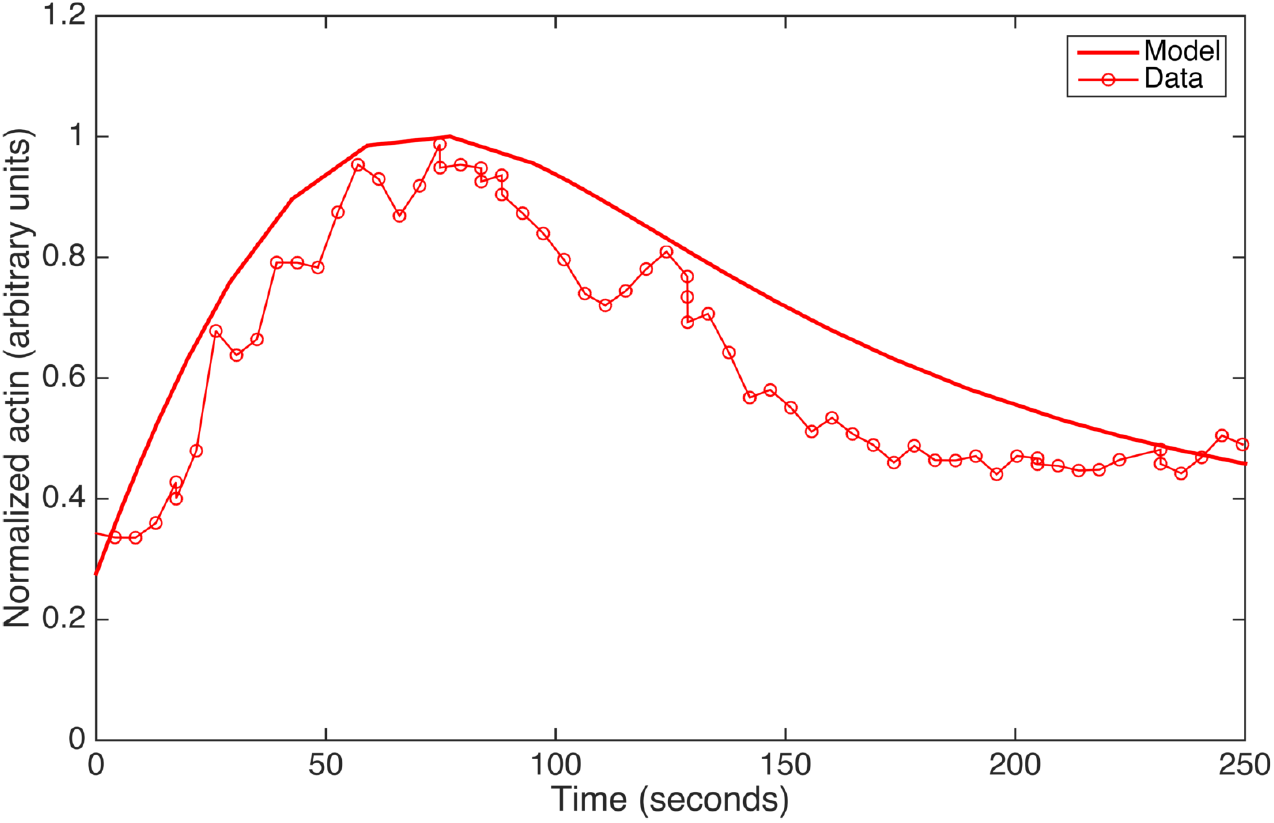
Plots showing the comparison of actin filament concentration between our model (solid line) simulated until 250 s and experimental data (dots), from Choi et al. [2008]. Data is adapted by permission from Macmillan Publishers Ltd: Nature Cell Biology. Choi et al, Nature Cell Biology, 10, 9 (2008), copyright 2008.

Experimental data shows the sequence of appearance where after actin appears in the position of adhesion in the lamellipodium prior to the initiation of nascent adhesion assembly. The time difference between the two assemblies is around 15 − 20 s. Our model solutions can quantitatively predict this time lag, as shown in Fig. 5, where in our model 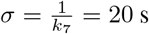.

**Figure 5:**
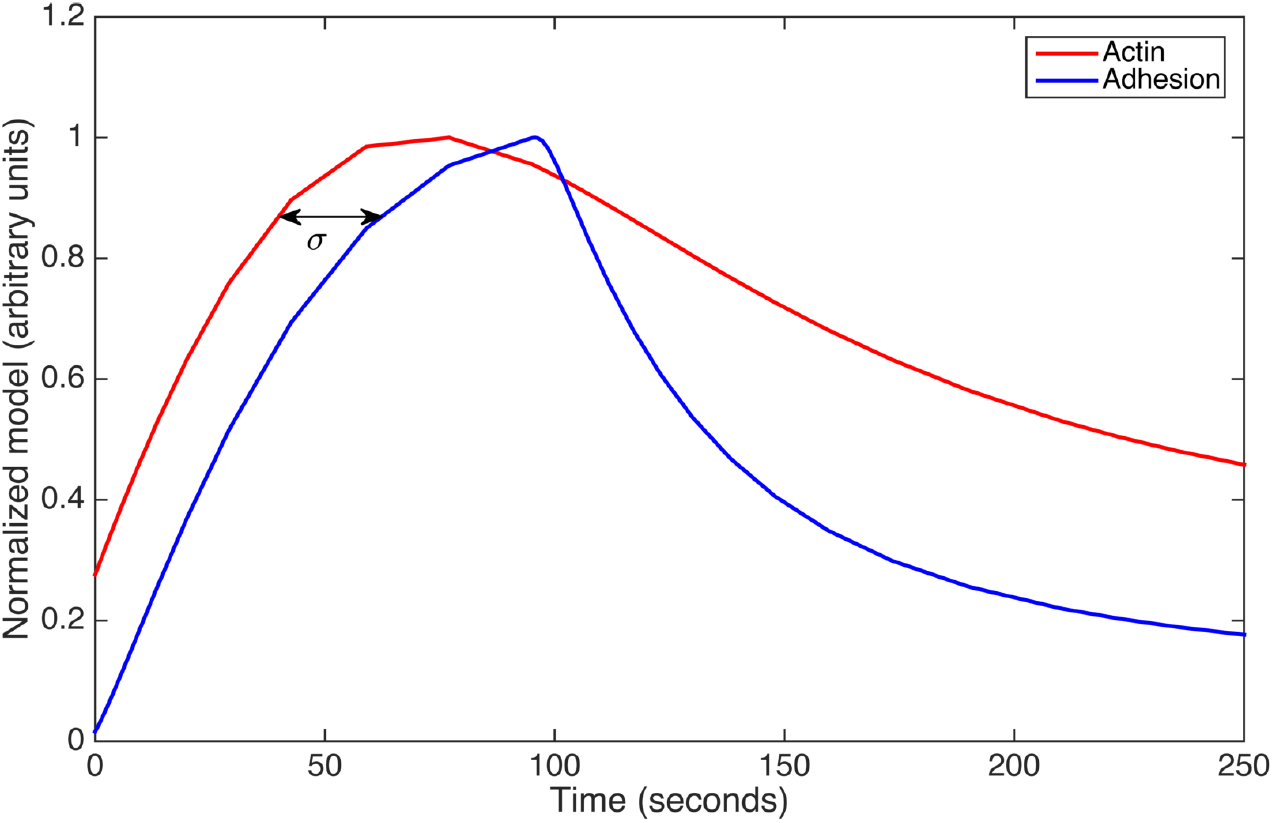
Plots showing model solutions of actin (red line) and adhesion (blue line) with time lag between actin and adhesion assemblies, represented by *σ*.

For the ECM displacement, the time delay *τ* = 95 s dictates that no displacement occurs until this point. During the initial period (0 *< t < τ*), forces generated by the assembling adhesion accumulate. After *τ* = 95 s, the effect of these forces on displacement becomes dominant, leading to an increase in displacement until it reaches a maximum, followed by a decrease as adhesions eventually disassemble. The solution depicted in Fig. 6 shows one cycle of ECM displacement corresponding to a single adhesion assembly and disassembly event. If subsequent nascent adhesions were to assemble before the current one completely disassembles, it would likely prevent the displacement from fully returning to its initial low value. The increasing displacement reflects the forces exerted on the ECM by mature nascent adhesions, which aligns with studies suggesting that nascent adhesions act as force generators that propel cell migration Beningo et al. [2001]. No experimental data for molecular ECM displacement at the scale considered in our model are currently available for direct comparison.

**Figure 6:**
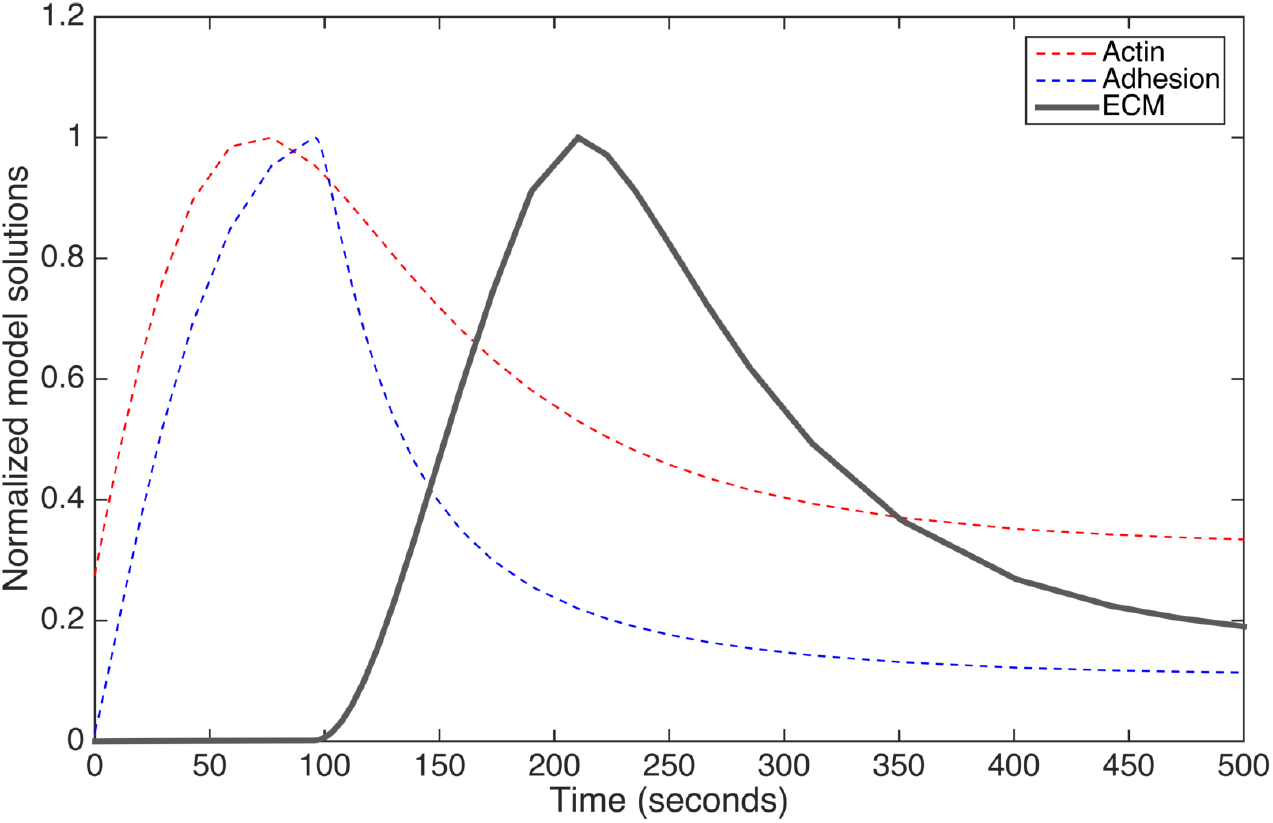
Plot showing model solution of ECM compartment containing integrin-ECM displacement using the parameters presented in Table 1 and with time delay *τ* = 95 s. There is no data available for comparison.

## 7 Conclusions and Discussion

In this paper, we have presented a compartmental model to predict the temporal dynamics of nascent adhesions by considering the interactions between the actin filament cytoskeleton (driven by barbed end dynamics), nascent adhesions themselves, and the local ECM displacement near the cell surface where integrins are bound. Nascent adhesions represent the initial adhesive structures formed during cell-matrix interactions, playing a crucial role in cell migration on both 2D substrates and within 3D environments. While the underlying dynamics of nascent adhesions are complex, the molecular components and mechanical principles governing their assembly and disassembly are relatively less intricate compared to larger, more mature adhesions like focal complexes, focal adhesions, and fibrillar adhesions. Consequently, our study focused on modeling nascent adhesion as a specific phenomenon, emphasizing its temporal profile. Our model successfully integrates the influence of both intracellular (actin) and extracellular (ECM) components in generating the characteristic three phases of nascent adhesion dynamics: assembly, transient stability, and disassembly. The solutions obtained from our relatively simple system of ordinary differential equations with a single time delay demonstrate a good fit with experimental data.

The mechanisms underlying nascent adhesion assembly have been the subject of considerable research, with two primary viewpoints emerging: (i) assembly driven by the binding of integrins to the ECM, and (ii) assembly driven by actin polymerization Parsons et al. [2010]. In this work, we focused on analyzing the second hypothesis as the primary driver of nascent adhesion assembly. To reproduce the small size and transient nature of nascent adhesions, which undergo disassembly after a specific timeframe (95 s in our model, based on experimental observations), our model suggests that the extracellular environment exerts a negative feedback mechanism on adhesion turnover. This extracellular component, represented in our model by ECM ligands binding to cell surface integrins, leads to bond displacement, generating forces/tension on the ECM and consequently influencing the stiffness experienced by nascent adhesions. Given the nature of compartmental modeling, which inherently abstracts molecular details, forces and stiffness are not explicitly modeled. However, ECM stiffness, whose impact on nascent adhesions has been experimentally validated, is inversely related to displacement Bruinsma [2005], approximated by:

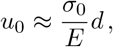

where *u*_0_ is molecular displacement, *σ*_0_ is the external stress applied to the adhesion, *d* is the adhesion size, and *E* is Young’s modulus, a measure of stiffness. The time delay introduced in the ECM displacement equation Eq. 4 can be interpreted as a relaxation time required for integrin-ECM bonds to respond to and act upon newly forming adhesion sites in their vicinity.

Our model effectively predicts the temporal dynamics of nascent adhesions, yielding results that are both qualitatively and quantitatively comparable to experimental data. Furthermore, our simulations provide insights into the significant contributions of both actin dynamics and ECM interactions in the formation and temporal profiling of these small, transient adhesions.

Despite its success in predicting and matching experimental observations, our simplified three-component model does not incorporate the intricate molecular details of intracellular and extracellular processes. Existing models have included more detailed representations of actin retrograde flow, barbed end dynamics, and myosin II activity for intracellular mechanisms Choi et al. [2008], Cirit et al. [2010], Shemesh et al. [2012], Welf et al. [2013], aspects that are not accounted for in our current formulation. Conversely, some models focus solely on the stiffness of the ECM interacting with integrins Peng et al. [2012], neglecting intracellular (or actin) dynamics. These limitations highlight opportunities for future theoretical studies to build upon our work by integrating more comprehensive intracellular and extracellular dynamics, recognizing that adhesion formation is indeed a bidirectional signaling process.

The findings from our model, along with those from previous studies, pose intriguing challenges for experimental investigations, particularly concerning the mechanisms of adhesion disassembly from an extracellular perspective. A deeper understanding of this aspect is crucial for elucidating the bidirectional signaling in cell-matrix adhesion. Gaining further insight into these processes may significantly contribute to a more complete understanding of complex cell migration events, such as those occurring during tumor invasion and metastasis.

